# Feeding ecology and ecological risks of the invasive fish *Coreoperca herzi* revealed by gut content DNA and environmental DNA metabarcoding

**DOI:** 10.64898/2026.03.20.713311

**Authors:** Satsuki Tsuji, Yusuke Hibino, Subaru Morimoto, Yugo Miuchi, Katsutoshi Watanabe

## Abstract

Understanding the dietary patterns of introduced predators is essential for assessing their impacts on freshwater ecosystems. Here, we investigated the feeding ecology of the invasive Korean perch (*Coreoperca herzi*) introduced to the Oyodo River system, Japan, by integrating gut content DNA metabarcoding and environmental DNA (eDNA) metabarcoding. Fifty specimens were collected, and prey taxa were identified using metabarcoding targeting fish, aquatic insects, and crustaceans. In parallel, eDNA metabarcoding of habitat water samples was used to assess prey availability and selectivity. The results revealed that the Korean perch prey extensively on aquatic insects and fish. Aquatic insect prey were dominated by epilithic clinger taxa inhabiting stone surfaces, particularly mayflies, suggesting visual-mediated prey selection. Fish predation was frequently detected even in small individuals (<100 mm SL), in contrast to previous studies based on conventional methods, indicating that piscivory begins early and ontogenetic dietary shifts are not pronounced. Furthermore, quantitative fish eDNA analysis showed a positive relationship between eDNA concentrations of prey species and predation frequency, indicating opportunistic feeding on abundant, size-accessible prey. By applying two metabarcoding approaches, this study provides an integrated assessment of prey utilisation and environmental context, highlighting ecological risks posed by the Korean perch to freshwater communities in Japan.

## Introduction

The introduction of non-native species can have serious impacts on biodiversity through competition with native species, predation, and pathogen transmission (Britton, 2023; Cucherousset and Olden, 2011; Mooney and Cleland, 2001). Among these, the effects of predation are particularly significant, potentially causing not only a reduction in prey populations but also widespread and irreversible ecosystem changes, via multiple ecological pathways, such as changes in behavioural and life history traits of species, shifts in interspecific relationships, and the collapse of community structures (Sih et al., 2010; Walsh et al., 2016). The extent and nature of these impacts largely depend on the predator’s dietary patterns; thus, elucidating the diet is essential for properly assessing the ecological impact of introduced predators (Dick et al., 2017; Wainright et al., 2021). Where dietary information is lacking, ecological impacts may be underestimated, leading to delayed countermeasures and erroneous management decisions.

The Korean perch (*Coreoperca herzi* Herzenstein, 1896; Fig. 1A), a carnivorous freshwater fish native to the Korean Peninsula, was first confirmed to have become established in Japan in 2017 in the Hagiwara River, a tributary of the Oyodo River system of Kyushu Island (Fig. 1B; Hibino et al., 2019). The Oyodo River is one of the major river systems in Kyushu Island, with a catchment area of 2,330 km^2^. The fish fauna of this river system is generally typical of western Japan(Saiki et al., 2023; https://www.nilim.go.jp/lab/fbg/ksnkankyo/); however, it is characterised by the presence of a loach species, *Cobitis sakahoko*. This species is endemic to this system and a single neighbouring river system (Nakajima and Suzawa, 2016; Oka et al., 2025). The population of the Korean perch has increased rapidly, with reports indicating its distribution expanded to multiple tributaries within the system by 2023, including the Okimizu River, a tributary located upstream of the Oyodo River No. 1 Dam (Fig. 1B; Hibino et al., 2022; Tsuji et al., 2024). As the Korean perch is a predator primarily feeding on aquatic insects and fish, it is considered to pose a high risk of predatory impacts and ecological competition for native species. Indeed, a marked decline in benthic fish populations has been reported in the Hagiwara River, where the perch was first recorded (Hibino et al., 2022). Additionally, quantitative fish environmental DNA (eDNA) surveys across the entire river system revealed the unexpectedly large-scale spread of the Korean perch and negative associations between the eDNA concentration of the perch and that of nine fish species, suggesting that predation by the perch may be causing a decline in native fish populations (Tsuji et al., 2024).

**Fig. 1.**
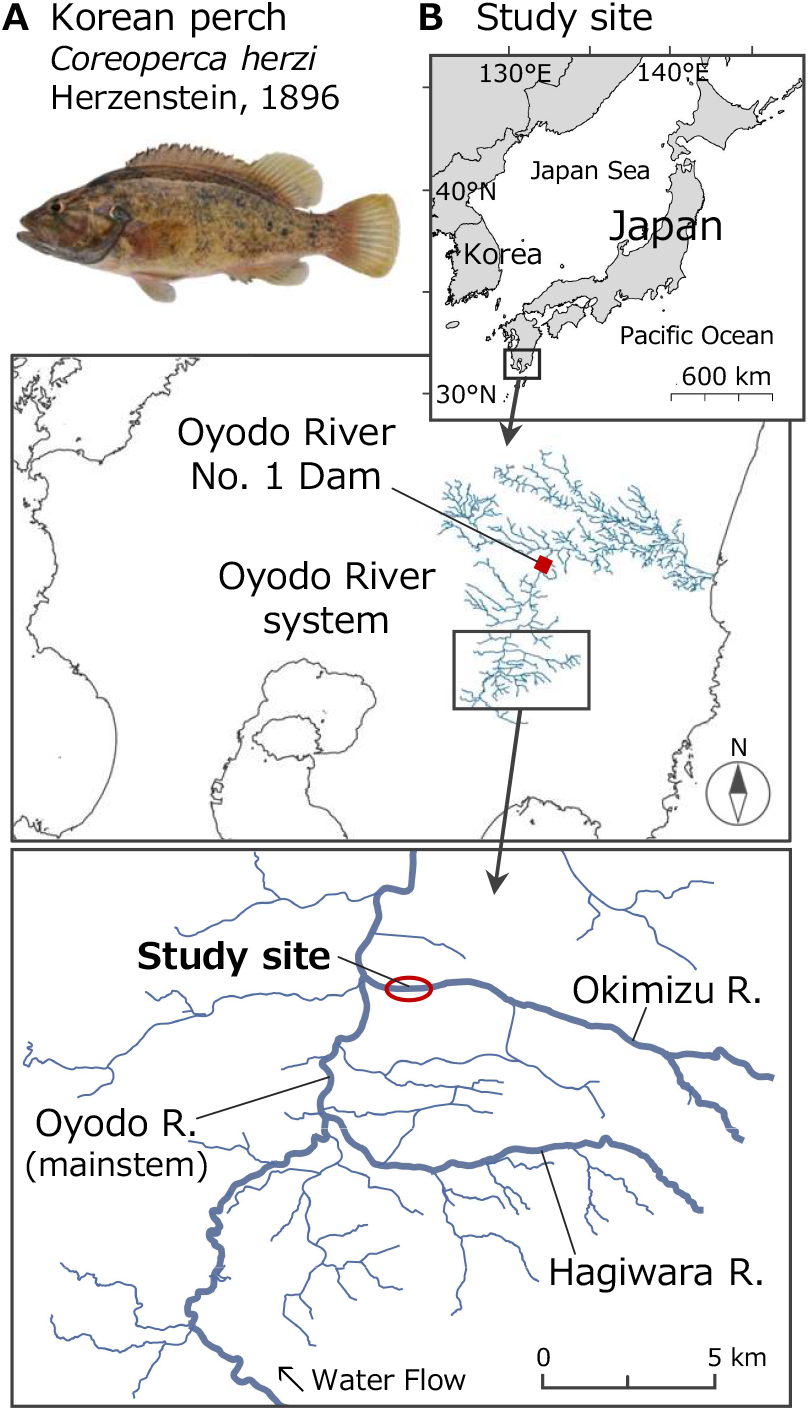
(A) The Korean perch, *Coreoperca herzi* Herzenstein, 1896; (B) overview of the study site

Three previous studies based on visual observations of stomach contents have reported that the Korean perch primarily preys on aquatic insects, particularly mayfly larvae (Ephemeroptera), as well as small fish, in the Hagiwara River (Akamine et al., 2025; Fujita and Yamanaka, 2025) and the Okimizu River (Akamine et al., 2025; Tsuji et al., 2024). However, numerous unidentified tissue fragments were also observed in these studies due to extensive digestion, indicating that the diversity of prey resources may have been underestimated. Furthermore, increases in body size during ontogeny frequently alter feeding selectivity in fishes (Duffy et al., 2010; Sánchez-Hernández et al., 2019a), and ontogenetic dietary shifts have been shown to influence both the functional roles of individuals and interspecific interactions within ecosystems (Sánchez-Hernández et al., 2019b). For the Korean perch, both Akamine et al. (2025) and Fujita and Yamanaka (2025) have suggested a dietary shift from insectivory to piscivory with growth, a pattern frequently observed in carnivorous fish (García-Berthou, 2002; Nolan and Britton, 2025; Shedd et al., 2015). However, because fish larvae are rapidly digested and difficult to detect visually, this may cause under-detection in stomach contents, particularly for smaller individuals that are more likely to consume them. Against this background, more detailed dietary data on the Korean perch are needed to reassess their potential ecological impacts.

To advance understanding of the dietary patterns of the Korean perch established in Japan, this study employed a combined approach incorporating conventional visual observation, gut content DNA metabarcoding, and eDNA metabarcoding. DNA metabarcoding approaches enable the identification of prey species from partially digested or morphologically unrecognisable gut contents, offering a more sensitive and better-resolved dietary assessment than visual inspection alone (Jakubavičiūtė et al., 2017; Pompanon et al., 2012). In addition, eDNA metabarcoding targeting fish, aquatic insects and crustaceans was used to estimate the availability of prey species at the study site, allowing assessment of whether the Korean perch exhibits selective predation on specific taxa or feeds opportunistically on abundant species. Specifically, this study aimed to (i) investigate dietary composition and (ii) examine ontogenetic dietary shifts through metabarcoding analysis of gut contents in the Korean perch, and (iii) infer prey selectivity for fish species through eDNA metabarcoding of water samples from the study site. This study provides a more accurate evaluation of the ecological impacts of the Korean perch on Japanese freshwater ecosystems and offers scientific insights that may inform future management and control strategies for this invasive species.

## Materials and Methods

### Field collection of Korean perch

In September 2024, 50 specimens of the Korean perch were collected from the lower reaches of the Okimizu River (approximately 500 m centred on 31.750056° N, 131.075417° E; Fig. 1B). This section was still at an early stage of Korean perch invasion and was selected as the survey site because it receives a continuous supply of food resources from the mainstem of the Oyodo River and upstream areas. The specimens were caught during night dives (after 10:00 PM) using a hand net (mesh size 5 mm), temporarily preserved in ice, and transported to the laboratory in a frozen state. For each specimen, the wet weight (± 1 g) and standard length (± 1 mm) were measured. Fulton’s condition factor (K) was calculated as K = 100 × W/SL^3^ (W = wet weight without gut contents, g; SL = standard length, cm). The individuals were dissected to determine sex and to extract the stomach and intestinal contents. The removed contents were separated into solids and semi-liquids and preserved in 99% ethanol.

### DNA extractions and library preparations for gut contents analysis

DNA extractions from the removed contents were performed using the DNeasy Blood & Tissue kit (Qiagen), following the manufacturer’s instructions. Solid samples were individually subjected to DNA extraction using either a portion or the entirety of the tissue following species identification through morphological observation. Semi-liquid samples were filtered using a coffee filter to collect the suspended particles. After drying at 56°C for 5 minutes to remove ethanol from the filter, DNA was extracted using the entire collected sample.

The DNA barcode regions of three taxonomic groups, i.e. fish, aquatic insects and crustaceans, were amplified in separate PCR reactions. For each target taxonomic group, the DNA region and primer set are as follows: fish, ca. 174 bp of 12S rRNA, MiFish-U primers (Miya et al., 2015); aquatic insects, ca. 200 bp of 16S rRNA, MtInsects-16S primers (Takenaka et al., 2023); decapod crustaceans, ca. 164 bp of 16S rRNA, MiDeca primers (Komai et al., 2019). All primers have been reported to have high detection rates in Japan across all target taxonomic groups. Details of the PCR conditions for each primer are provided in the supplementary file. Sequence analysis was outsourced to Novogene’s gigabyte-scale sequencing service, and 150 bp (fish and decapod crustaceans) or 250 bp (aquatic insects) paired-end sequencing was performed using Illumina NovaSeq X Plus.

### Environmental DNA analysis

To understand the fauna (fish, aquatic insects, crustaceans) at the capture site of the Korean perch, eDNA analysis using water samples was conducted. The water samples were collected in May 2024 for a different study. One litre of surface water was collected using disposable cups, immediately filtered through a GF/F filter, and the filter was frozen with dry ice. Water sampling was carried out during daylight hours. In the laboratory, eDNA was extracted from the filters using a spin column and DNeasy Blood & Tissue kit according to Tsuji et al. (2024). DNA library preparation was performed using MiFish-U primers, MtInsects-16S primers and MiDeca primers. In preparing fish eDNA libraries using MiFish-U primers, quantitative fish eDNA metabarcoding was performed by adding three types of standard DNA at known different concentrations (5, 25, 50 copies) to each primary PCR reaction (Ushio et al., 2018). These standard DNAs were amplified simultaneously with fish eDNA using the MiFish-U primers, enabling quantification of the number of DNA copies for each detected species. The standard sequences were developed by Tsuji et al. (2022). Details are shown in the supplementary file. Sequence analysis was outsourced to Novogene’s gigabyte-scale sequencing service, together with gut contents analysis.

### Bioinformatic analysis

The bioinformatics analysis was performed using the PMiFish pipeline ver. 2.4.1 (https://github.com/rogotoh/PMiFish; Miya et al., 2020). Briefly, the PMiFish pipeline includes the following steps: removal of low-quality tail reads, merging of the paired-end reads, removal of primer sequences, quality filtering, removal of low-frequency reads, denoising by UNOISE and species assignment. The details of the parameter settings for each analysis step are shown in the supplementary file. For the species assignment step, the sequence data of each target region were batch downloaded from NCBI to construct the reference database. For fish, the sequences of three standard DNAs were added to the reference database.

For the quantitative fish eDNA metabarcoding, the conversion of the number of sequence reads for each species to DNA copy number was performed per sample according to Ushio et al. (2018). To obtain a sample-specific standard line, linear regression analysis (lm function in R software ver. 4.5.2; R Core Team, 2025) was performed using the sequence reads of standard DNAs and their known copy numbers, with the intercept set to zero. For each sample, the number of eDNA copies per litre for each detected species was calculated using the following equation: qMiSeq eDNA concentration = (the number of sequence reads/regression slope of the sample-specific standard line) × 100. The detection of saltwater fish species was considered to be due to contamination from domestic wastewater, and they were removed from the data because the survey site was in freshwater.

### Gut content analysis and statistical analyses

Because quantitative comparisons of individual prey items in gut contents are difficult using the metabarcoding approach, only the predation frequency of each prey species (i.e. occurrence rate among individuals; %F) was calculated using the following formula:

%F = [number of specimens containing a prey species / (total number of specimens – number of individuals with empty guts)] × 100

All statistical analyses and graphic illustrations were carried out using R software, and statistical values were evaluated at a significance level of α = 0.05. The relationship between SL (mm) and wet weight (without gut contents, g) or Fulton’s condition factor K was examined using Spearman’s rank correlation test.

Differences in diet composition of each specimen were visualised using the non-metric multi-dimensional scaling (NMDS) method with 999 separate runs of the Jaccard similarity matrix using the metaMDS function in the vegan package ver. 2.5.6 in R. The Jaccard similarity matrix was calculated from the binary presence–absence data of detected taxa by gut contents DNA metabarcoding. The NMDS stress was calculated to assess the adequacy of the ordination. Additionally, to evaluate the significance of the effect of specimen body size on the diet composition, permutational multivariate analysis of variance (PERMANOVA) was conducted with 999 permutations using the adonis2 function in the vegan package. To examine whether ontogenetic dietary shifts towards piscivory occurred, we modelled the presence or absence of fish predation (binary response: 1 = fish detected, 0 = not detected) as a function of standard length (SL, mm) using a generalised linear model (GLM) with a binomial error distribution and logit link function.

Furthermore, to examine whether the Korean perch exhibited selectivity among prey species, the relationship between the eDNA concentration of fish species at the survey sites and their %F was analysed. Typically, eDNA concentrations reflect the biomass of target species (Rourke et al., 2022; Tsuji et al., 2022a). However, species that are presumed to grow to a size (total length, >20 cm) where they cannot be physically preyed upon by the Korean perch were excluded from the analysis, as the presence of large individuals could introduce noise into the estimation of the relationship between eDNA concentration and predation (see Discussion). A GLM was fitted with %F of each species as the response variable and the eDNA concentration of the same species as the explanatory variable, using a Poisson error distribution with a log-link function. In addition, the Korean perch were simply divided into two size groups (58–99 mm SL, small, *n* = 21; 100–165 mm SL, large, *n* = 29), and %F was calculated for each group. Differences between size groups were evaluated using Fisher’s exact test to compare the degree of piscivory associated with growth.

## Results

The 50 captured specimens of the Korean perch ranged from 58 to 165 mm in standard length (SL) (104 ± 22 mm, average ± SD; Table S1), showing a strong positive correlation between SL and wet weight (Spearman’s rank correlation test, rho = 0.98, *p* < 0.001; Fig. S1). Fulton’s condition factor (K) ranged from 2.5 to 3.8 (3.1 ± 0.3) with no significant relationship to SL (Spearman’s rank correlation test, rho = 0.05, *p* = 0.75). The empty gut rate was 10% (5 of 50 specimens), with one in the small group and four in the large group (Fig. S1). Details of prey organisms detected from each specimen are shown in Tables 1 and S2.

**Table 1.** Gut contents of the Korean perch (*n* = 45, excluding individuals with empty guts) as detected by visual observation and metabarcoding. Occurrence rate of gut contents (%F) was calculated as (number of specimens feeding on a given prey species / 45 individuals) × 100.

| Group |  | Species<br>(sp./spp./family) | Number of specimens<br>feeding on each prey<br>species | Detected method |  | Occurrence<br>rate of the gut<br>contents (%F) |
| --- | --- | --- | --- | --- | --- | --- |
|  |  |  |  | Visual &<br>Metabarcoding | Metabarcoding<br>only |  |
| Fish | Pelagic fish | *Nipponocypris temminckii* | 1 | 1 | 0 | 2.2 |
|  |  | *Gnathopogon elongatus* | 4 | 2 | 2 | 8.9 |
|  |  | *Opsariichthys platypus* | 18 | 9 | 9 | 40.0 |
|  |  | *Rhynchocypris oxycephalus jouyi* | 3 | 0 | 3 | 6.7 |
|  | Benthic fish | *Misgurnus anguillicaudatus* | 3 | 0 | 3 | 6.7 |
|  |  | *Odontobutis obscurus* | 2 | 0 | 2 | 4.4 |
|  |  | *Pseudogobio esocinus* | 22 | 12 | 10 | 48.9 |
|  |  | *Rhinogobius* sp. | 3 | 1 | 2 | 6.7 |
| Shrimp |  | *Neocaridina denticulata* | 4 | 0 | 4 | 8.9 |
|  |  | *Neocaridina* spp. | 3 | 2 | 1 | 6.7 |
| Aquatic insects | Dragonfly | Corduliidae | 2 | 0 | 2 | 4.4 |
|  |  | Gomphidae | 5 | 2 | 3 | 11.1 |
|  | Stonefly | Perlidae | 1 | 0 | 1 | 2.2 |
|  |  | Nemouridae | 1 | 0 | 1 | 2.2 |
|  | Beetles | Psephenidae | 1 | 0 | 1 | 2.2 |
|  | Mayfly | Ephemerellidae | 15 | 0 | 15 | 33.3 |
|  |  | Caenidae | 1 | 0 | 1 | 2.2 |
|  |  | Heptageniidae | 37 | 2 | 35 | 82.2 |
|  |  | Potamanthidae | 21 | 0 | 21 | 46.7 |
|  |  | Baetidae | 30 | 2 | 28 | 66.7 |

Solid gut contents were obtained from 17 specimens (34%) and classified into three families of aquatic insects, five fish species, and one shrimp species through individual morphological observations and DNA barcoding (Table S2). Semi-liquid contents were obtained from 45 specimens (90%). Metabarcoding of each specimen detected 10 families of aquatic insects, eight fish species, and two shrimp species, including all species identified in the solid contents of any of the specimens examined (Table S2). The occurrence rate of the gut contents (%F) was highest for aquatic insects (98.0%, 44 specimens), fish (66.7%, 30 specimens) and shrimp (17.8%, four specimens) (Fig. 2). Twenty-seven specimens (60.0%) preyed on both aquatic insects and fish, while two specimens (4.4%) preyed on both aquatic insects and shrimp, and another two (4.4%) preyed on all three taxonomic groups. In addition to the prey organisms obtained from gut contents, partially digested *Pseudogobio esocinus* (approximately 65 mm SL) and *Opsariichthys platypus* (approximately 35 mm SL) were confirmed to have been regurgitated into the container during specimen transport.

**Fig. 2.**
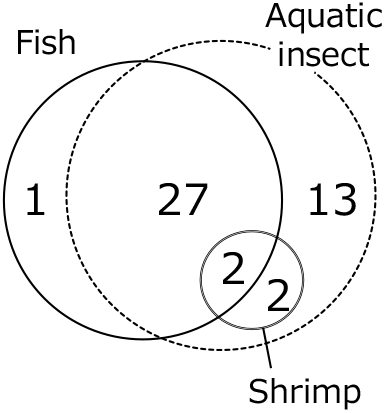
Number of Korean perch specimens that preyed on each taxonomic group (*n* = 45, excluding individuals with empty stomachs)

Aquatic insects were mostly limited to epilithic clinger taxa associated with stones and other substrates, and most of them were mayflies (e.g. Heptageniidae, Baetidae; Tables 1, S2). Among fishes, %F was higher for two species: *P. esocinus* (48.9) and *O. platypus* (40.0), while other species were rarely preyed upon (≤8.9). When separated into pelagic and benthic fish, %F was 57.8 and 66.7, respectively.

Quantitative fish eDNA metabarcoding revealed 19 taxa (mostly species), including the Korean perch, at the survey site (Table S3). Among these, the top 10 species in descending order of eDNA concentration were: *Opsariichthys platypus*\*, *Cyprinus carpio, Carassius* sp., *Carassius cuvieri, Nipponocypris temminckii*\*, *Rhinogobius* sp.*, *Pseudogobio esocinus*\* (*: adult fish TL ≤ 20 cm). A GLM analysis conducted to examine prey preference revealed a significant positive relationship between the eDNA concentration of each fish species (adult fish TL ≤ 20 cm) and %F of the species (Table S3; GLM, *p* < 0.001, Fig. 3), suggesting largely abundance-dependent predation. Furthermore, although no quantitative assessment was performed, eDNA metabarcoding of aquatic insects revealed high read frequencies of mayflies (1.95–42.9%) and stoneflies (0.11–1.95%), which were frequently preyed upon by the Korean perch (Table S4). Contrastingly, although black fly larvae (Simuliidae) were not detected in gut content metabarcoding, their read frequency in eDNA metabarcoding was relatively high (6.6%).

**Fig. 3.**
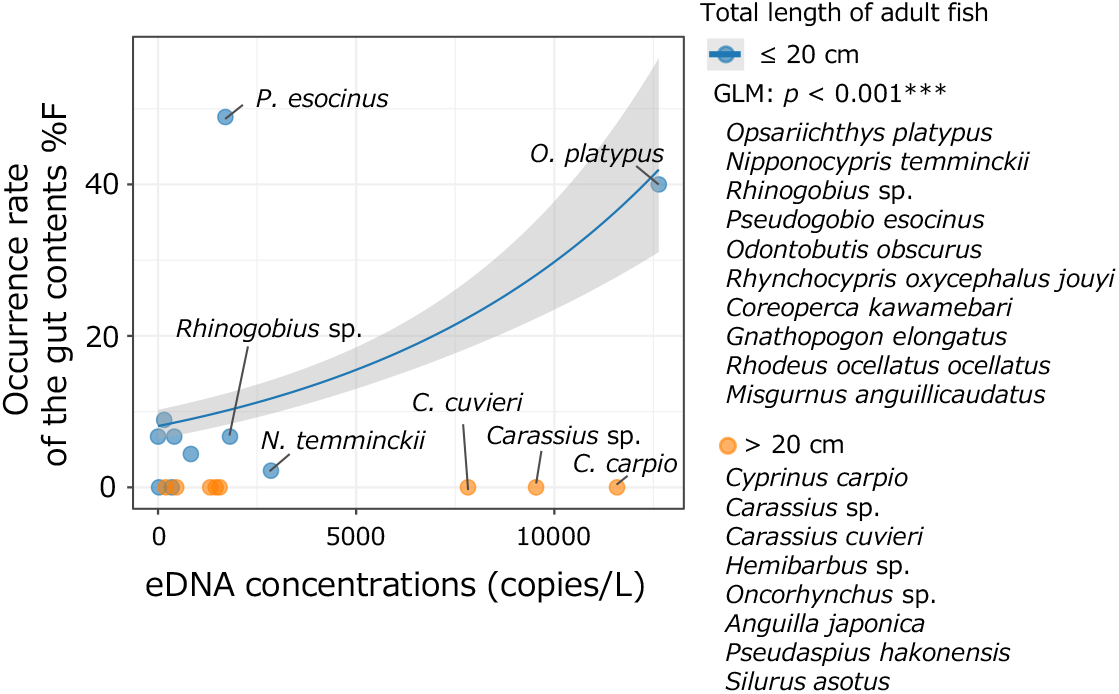
Relationship between the eDNA concentration of each fish species in the water sample from the study site and their occurrence frequency in the gut contents (%F). For species with an adult total length of ≤20 cm, GLM analysis indicated a significant positive relationship (*p* < 0.001)

The relationship between each specimen’s SL and prey composition was examined using NMDS and PERMANOVA (stress = 0.16, Fig. 4A). No significant relationship was found when using binary data for each prey species (PERMANOVA, *p* = 0.25). The GLM analysis also indicated no significant relationship between the probability of fish predation and body size of the Korean perch (*p* = 0.25; Fig. 4B). The small group of the Korean perch exhibited a %F of 70.0 for fish (*n* = 20), whereas the large group showed a %F of 64.0 (*n* = 25) (Table S2), indicating no significant difference between the two groups (Fisher’s exact test, *p* = 0.76).

**Fig. 4.**
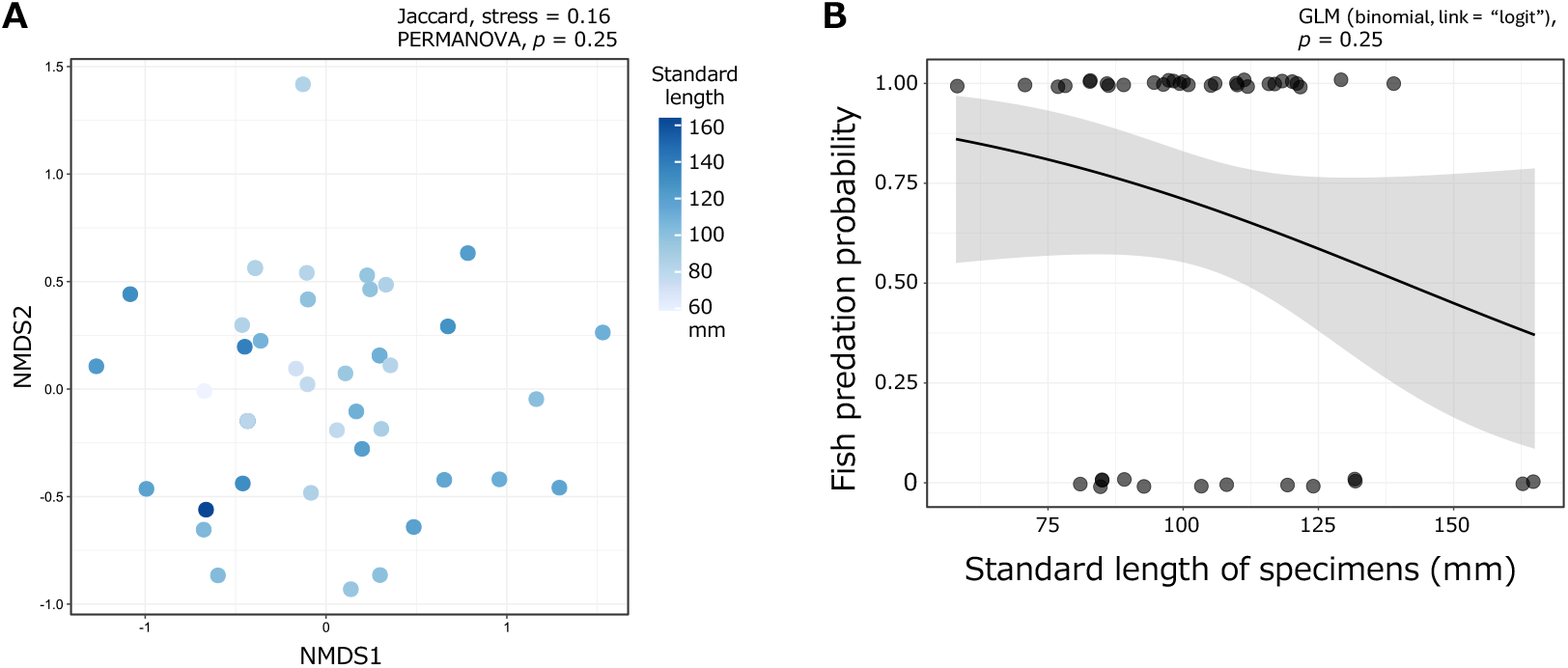
(A) NMDS ordination plot for prey composition based on binary gut content data, with each specimen’s standard length indicated by colour gradients. The NMDS stress value was 0.16. No significant relationship was detected between specimen standard length and prey composition (*p* = 0.25). (B) Presence or absence of fish predation by each specimen plotted as a function of standard length. Solid lines indicate the estimated probability of fish predation (GLM, *p* = 0.25). Plots are vertically staggered to minimise overlap. Shaded areas in both panels represent 95% confidence intervals

## Discussion

This study provides new insights into the feeding ecology of the Korean perch introduced into the Oyodo River, Japan. Notably, in contrast to previous reports (Akamine et al., 2025; Fujita and Yamanaka, 2025), predation of fish was detected even in relatively small individuals, indicating that fish consumption begins at an early stage of growth. In the following sections, these findings are discussed in the context of prey availability, behavioural flexibility, and potential ecological impacts on native freshwater communities. Finally, the advantages and limitations of combining gut content DNA metabarcoding with eDNA metabarcoding are discussed to clarify the methodological implications for future assessments of invasive predator–prey interactions.

### Dietary composition of the Korean perch

This study revealed that the introduced Korean perch, established in the Oyodo River system, Japan, actively preys on both aquatic insects and fish. This finding is consistent with previous studies based on visual observations of stomach contents (Akamine et al., 2025; Fujita and Yamanaka, 2025; Tsuji et al., 2024), suggesting that this species may exert strong predatory pressure on local freshwater ecosystems. Furthermore, despite the high diversity of aquatic insects detected by eDNA analysis in the Okimizu River, the Korean perch appeared to preferentially prey on epilithic taxa, such as mayflies and stoneflies, that actively crawl on submerged stone surfaces. This prey selectivity suggests that the perch primarily relies on vision to search for prey.

Compared with our observations in the Okimizu River, in the native range of the species in South Korea, as well as in the Hagiwara River, where the Korean perch was first recorded in the Oyodo River system, the species has been reported to prey on a broader range of aquatic insect types, including black fly larvae (Simuliidae), which are epilithic, attached benthic insects that adhere to stone surfaces via silk-like secretions (Byeon, 2017; Akamine et al., 2025; Fujita & Yamanaka, 2025). In the Okimizu River, however, black fly larvae were not detected in the stomach contents, despite being common in the environment. The differences in diet between the survey sites may reflect differences in prey availability and/or the population density of the perch. In the Hagiwara River, the population density of the Korean perch has already increased to a high level, and prey depletion has been noted (Hibino et al., unpublished), suggesting that foraging by the perch may have been constrained. When prey organisms are abundant, the Korean perch may selectively target particular taxa; however, under prey scarcity, they may broaden their diet to include substrate-attached (e.g. Simuliidae) or sediment-dwelling taxa (e.g. Chironomidae) (Byeon, 2017; Akamine et al., 2025; Fujita & Yamanaka, 2025). Such feeding and behavioural flexibility suggests that, as populations increase, predation pressure may expand across a wider range of aquatic insect groups. Given the central ecological roles of aquatic insects, including their contribution to nutrient cycling and trophic linkages between aquatic and terrestrial ecosystems (Nakano and Murakami, 2001), predation by the Korean perch could alter community structure and ecosystem functioning in introduced habitats.

This study suggests that the Korean perch frequently preys not only on benthic fish but also on small pelagic fish. Generally, benthic fish are considered easy prey for the Korean perch due to their limited swimming ability. Indeed, Hibino et al. (2022) reported a marked decline in benthic fish populations in the Hagiwara River, following the establishment and increase in the population density of the Korean perch, based on underwater visual surveys. On the other hand, while no clear reduction in biomass was reported for pelagic fish through visual surveys, the pronounced lure-chasing behaviour exhibited by the Korean perch has raised serious concerns regarding its capacity to prey effectively upon pelagic fish (Hibino et al., 2022). The results of the present study provide empirical support for this concern. Furthermore, comparisons of the body sizes of prey items identified in this study with those of Korean perch specimens indicated that the perch is capable of preying on fish up to approximately half its own body size. For example, the largest recovered prey item (*Pseudogobio esocinus*, 65 mm SL), although not assignable to a specific predator specimen due to regurgitation during transport, would correspond to approximately 40% of the body size of the largest perch specimen (165 mm SL) and 63% of that of a median-sized perch specimen (104 mm SL). The Korean perch is known to grow to a maximum total length of 300 mm (approximately 240–270 mm SL) (Uchida, 1935). Given this growth potential, fish species reaching adult sizes of around 200 mm in total length may face predation risk from the Korean perch at any growth stage. Furthermore, not only small□bodied species but also medium□to large□bodied species may face long□term impacts such as reproductive suppression and consequent population decline.

Regarding fish, no clear prey preference was detected in the Korean perch at the study site; it likely feeds indiscriminately on species that fall within a vulnerable size range and are abundant in the community (Fig. 3). For example, the pelagic fish *Opsariichthys platypus*, which had a high %F and eDNA concentration in this study, was the most abundant species in the survey area (as observed by Y.H.), suggesting a relatively high encounter frequency with the Korean perch. As even adult *O. platypus* fall within the size range that the Korean perch can prey on, they are likely to have been frequently utilised as a major food resource. Similarly, the benthic fish *P. esocinus*, which also exhibited a high %F, was frequently observed at night (as observed by Y.H.), although eDNA concentrations were likely underestimated due to its burrowing behaviour in sand during daylight hours, when the water samples were collected. In contrast, *Nipponocypris temminckii* exhibited a relatively low %F despite relatively high eDNA concentrations. This species favours slower-flowing habitats, which may reduce its encounter probability with the Korean perch, which prefers flowing water and riverbed interstices. These findings indicate that fish predation by the Korean perch extends broadly to species that are accessible in terms of size and population density, even if some degree of species-level selectivity is present.

This study and previous studies consistently suggest that predation of crustaceans by the Korean perch is generally infrequent; however, this finding does not necessarily imply that the perch exerts low predation pressure on crustaceans. In the Okimizu River and the Hagiwara River, the sites examined to date, crustacean populations were inherently low, and crustaceans were scarcely observed during collection surveys. Thus, the low representation of crustaceans in the diet of the Korean perch likely reflects the limited availability of this prey group in the local environment. Should future invasions of the Korean perch occur in water systems with abundant crustaceans, it will be necessary to reassess the predation pressure on crustaceans and its ecological consequences.

### Dietary patterns during ontogeny

Previous studies suggested that the fish-eating habits of the Korean perch may increase with its growth (Akamine et al., 2025; Fujita and Yamanaka, 2025). In contrast, this study found no significant relationship between body size and the composition of gut contents or the presence of fish consumption. Our sampling did not include individuals smaller than 58 mm SL, whereas previous studies incorporated smaller size classes (≥27 mm SL; Table S5). Nonetheless, even presumed age□0 individuals were found to have preyed on not only aquatic insects but also small fish.

The present result implies that the Korean perch can prey on fish from a relatively early life stage, suggesting that ontogenetic dietary shifts are not pronounced. Since this study employed DNA metabarcoding, dietary evaluation was necessarily based solely on occurrence frequency (%F), and a direct comparison with previous studies that used %IRI, accounting for occurrence frequency, numerical percentage and weight percentage (Akamine et al., 2025; Fujita and Yamanaka, 2025; Pinkas et al., 1970), is therefore limited. Nevertheless, DNA metabarcoding has high sensitivity for detecting partially digested gut contents and minute prey items, and thus may reduce the underestimation of fish predation. Indeed, in our study, fish were visually detected in only a subset of specimens for which fish predation was indicated by metabarcoding (4 of 14 specimens, 29%, in the <100 mm SL group; 9 of 16 specimens, 56%, in the ≥100 mm SL group; Table S2). Although differences in sampling periods [September 2024 in this study vs. March–November 2023 in Akamine et al. (2025) and September 2023 in Fujita & Yamanaka (2025)] should be taken into account, the low reliance on fish prey in small Korean perch individuals reported in previous studies may have been underestimated. In this study, fish predation was detected in 70.0% of individuals in the 58–99 mm SL group, which did not significantly differ from the value (64.0%) in the 100–165 mm SL group. These values were markedly higher than those reported by Fujita & Yamanaka (2025) from the Hagiwara River (%F, 1.9% for 27–79 mm SL and 18.2% for 80–133 mm SL). Although these occurrence frequency data are not directly comparable with the %IRI values, Akamine et al. (2025) reported %IRI values of <0.5% in the Hagiwara River and 16.3% in the Okimizu River for individuals 60–119 mm SL, markedly lower than those for individuals ≥120 mm SL (69.9% and 88.8%, respectively), a pattern that contrasts with the results of the present study. In future studies, it would be desirable to evaluate the diet and ecological impacts of the Korean perch by integrating visual observations with DNA metabarcoding, while also including smaller individuals in the sampling regime and considering contexts that differ in invasion stage, population size, and faunal composition.

### Gut content DNA metabarcoding: limitations and complementary insights from eDNA

By combining gut content DNA metabarcoding with visual observation, this study conducted a highly sensitive assessment of the dietary patterns of the Korean perch, markedly enhancing our understanding of its dietary profile. However, DNA metabarcoding has several technical limitations (Pompanon et al., 2012). Specifically, the number of sequence reads does not necessarily correlate with prey abundance or biomass, making it impossible to calculate quantitative indices such as the proportions of the stomach contents by weight (%W) or the percentage index of relative importance (%IRI), which have been widely used in conventional dietary studies. Furthermore, contamination of the gut sample with the predator’s own DNA or the DNA of the prey’s prey can make it difficult to assess the occurrence of cannibalism or secondary predation (Sheppard et al., 2005). In this study, DNA from the Korean perch was frequently detected; however, no recognisable perch bodies were visually observed in the gut contents, and it therefore remains unclear whether the detected DNA originated from the predator itself or from conspecific prey that had been digested. Meanwhile, the only potential piscivorous species identified was *Odontobutis obscurus*, which occurred at low frequency in the gut contents (Table 1; Hosoya, 2019), suggesting that, at least for fish, overestimation of predation rates due to secondary predation is likely minimal.

This study incorporated eDNA metabarcoding to assess the availability of prey taxa, thereby complementing gut contents DNA metabarcoding results and offering additional insights into predation patterns. In particular, quantitative eDNA metabarcoding with MiFish primers helps to infer the feeding behaviour of the Korean perch, including whether it selectively targets particular taxa or opportunistically consumes abundant taxa. Investigating prey community composition using conventional capture or observation methods requires considerable effort and time, whereas eDNA analysis enables a simpler, simultaneous investigation of multiple taxa. This highlights the value of integrating gut content DNA metabarcoding and eDNA metabarcoding in elucidating prey selection patterns and in understanding how prey resource utilisation is related to the environmental context.

## Data availability

All raw sequences and metadata are deposited in the DDBJ Sequence Read Archive with the accession numbers: DRR914636–DRR914803 (satsuki_ddbj-0019_Run_0001-0168). The correspondence between each specimen ID and its filename is listed in Table S6.

## Acknowledgements

This study was supported by JSPS KAKENHI Grant Number 23K13967.

## Author Contributions

Study conception and design: Satsuki Tsuji, Yusuke Hibino, and Katsutoshi Watanabe; Field sampling: Satsuki Tsuji (eDNA sampling), Yusuke Hibino (fish collection); Fish dissection and gut content sampling: Satsuki Tsuji, Subaru Morimoto, Yugo Miuchi, Katsutoshi Watanabe; Molecular experiments and data analysis: Satsuki Tsuji; Writing of the paper: Satsuki Tsuji wrote the first draft and finalised the manuscript with substantial input from all authors.

## Compliance with Ethical Standards

All experiments were performed with attention to animal welfare. Based on current laws and guidelines in Japan relating to animal experiments, the collection of fish and the use of DNA samples are permitted without requiring any ethical approvals from authorities.

## Figures

Fig. S1. Relationships between specimen standard length and wet weight (excluding gut contents), and Fulton’s condition factor (K). The shape of each point indicates the sex of the specimen. The lower panel shows the frequency distribution of standard length

