## Supplementary figures and images for "Feeding ecology and ecological risks of the invasive fish *Coreoperca herzi* revealed by gut content DNA and environmental DNA metabarcoding"

### Supplemental figure

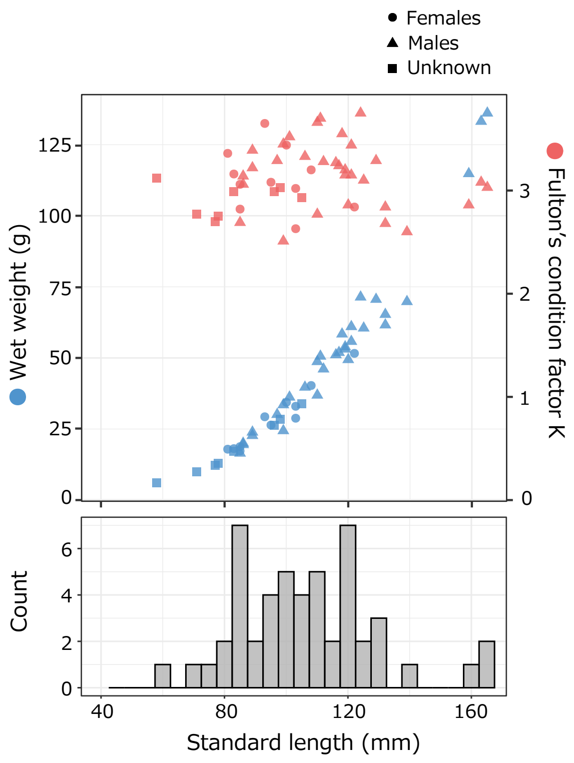
